# Establishment of the mayfly *Cloeon dipterum* as a new model system to investigate insect evolution

**DOI:** 10.1101/494674

**Authors:** Isabel Almudi, Carlos Martín-Blanco, Isabel M. García-Fernandez, Adrián López-Catalina, Kristofer Davie, Stein Aerts, Fernando Casares

**Affiliations:** GEM-DMC2 Unit, the CABD (CSIC-UPO-JA). Ctra. de Utrera km1 41013,Sevilla SPAIN; Laboratory of Computational Biology, VIB Center for Brain & Disease Research, Herestraat 49, 3000 Leuven, Belgium; Department of Human Genetics, KU Leuven, Oude Markt 13, 3000 Leuven, Belgium

## Abstract

The great capability of insects to adapt to new environments promoted their extraordinary diversification, resulting in the group of Metazoa with the largest number of species distributed worldwide. To understand this enormous diversity, it is essential to investigate lineages that would allow the reconstruction of the early events in the evolution of insects. However, research on insect ecology, physiology, development and evolution has mostly focused on few well-established model species. The key phylogenetic position of mayflies within Paleoptera, as the sister group of the rest of winged insects and life history traits of mayflies make them an essential order to understand insect evolution. Here, we describe the established of a continuous culture system of the mayfly *Cloeon dipterum* and a series of experimental protocols and -omics resources that allow the study of its development and its great regenerative capability. Thus, the establishment of *Cloeon* as an experimental platform paves the way to understand genomic and morphogenetic events that occurred at the origin of winged insects.

## Introduction

Insects are the most diverse group of Metazoa, harbouring the largest number of animal species [1]. Insects comprise more than thirty extant orders distributed worldwide - they are found in all sorts of habitats with the exception of marine environments [2]. Despite the fact that other animals populated the land before insects, like chelicerates and myriapods [3–5], the appearance of winged insects meant a complete biological revolution with profound effects on the history of life on earth. The colonization of the air allowed insects unprecedented dispersal capacities and novel ecological interactions-such as their role as pollinator agents that drove the further coevolution of insects and angiosperms.

Although the impact that the appearance of insects had in the shaping and evolution of, not only their own group, but also other phyla and even kingdoms, our knowledge of insects comes mainly from work on a handful of well-established model species. Among them, *Drosophila melanogaster*, which is one of the best-studied model organisms, broadly used in multiple fields of research, including the evo-devo field [6–8]. Probably the second more used insect in evolutionary and developmental studies is *Tribolium castaneum* (Coleoptera), followed by some butterflies and moths (Lepidoptera) species. In addition to these established models, other dipterans with important impact in human health (as vectors transmitting diseases: *Anopheles, Glossina, Aedes*) and economy (becoming agricultural pests: *Ceratitis capitata, D. suzukii*) have been studied in more detail. Unfortunately, these insect orders are all part of the holometabola group of hexapoda, which appeared relatively recently within the insect phylogeny ([9] and references therein). Some efforts have been made in order to fill this gap of studies in hemimetabola animal systems, such as *Oncopeltus fasciatus* [10], *Blattella germanica* [11] and water striders [12, 13].

This dearth of laboratory models is even more acute in the case of early branching groups of insects that correspond to the first representatives of the crucial biological and ecological transitions mentioned above. Such transitions are for instance, key adaptations to terrestrial life such as the development of the extraembryonic tissue amnion and serosa [14–18], the establishment of early embryo segmentation mechanisms and the transition from short-to long-germ band embryos [19–23], the basal organization of the head [24, 25], or the origin of wings and the capacity to fly (an issue that is currently hotly debated [26–34]). Overall, what these examples reveal ultimately is the need of developing and establishing new model systems, in particular around the nodes of the tree where these key novelties originated.

The advent of new technologies that permit the editing of genomes and especially the appearance of next-generation sequencing (NGS) techniques allows the re-examination of long-standing questions in Evolutionary Biology using comparative approaches. For instance, the use of functional genomics methods and other “-omic” techniques enables the interrogation of genomes and transcriptomes in search for changes in gene content (appearance of new lineage-specific gene families) and regulatory elements that promoted the origin and evolution of these key innovations that appeared for the first time in insect lineages. However, one of the challenges that researchers are encountering is getting access to the biological material, especially at the desired developmental stage for a particular study. Thus, there is a great interest in increasing the number of emergent model organisms that, due to their key phylogenetic position or their specific traits, would permit evo-devo studies in the precise clade of interest. Here, to contribute in this direction, we developed an Ephemeroptera laboratory model, *Cloeon dipterum*.

Ephemeroptera (mayflies) is an order of winged hemimetabola insects that live in freshwater ecosystems. The Ephemeroptera order has over 3000 species distributed in 40 different families approximately [35, 36]. Mayflies belong to an ancient group of insects that were present already in the late Carboniferous or early Permian period [1]. Mayflies have a life cycle that consists of two well-defined phases. The aquatic phase, that comprises embryogenesis and nymphal stages and the terrestrial phase, which consists on a sexually immature subimago and a sexually active imago (Fig 1). Their aquatic phase makes mayflies ideal as bioindicators of the quality of freshwater ecosystems [37–39], while their terrestrial phase contributes to population dispersal; thus, mayflies have been used to investigate biogeographical events, such as dispersion and colonization of new communities [40–42]. The embryogenesis occurs in a variable amount of time that can range from days until months, depending on the species and environmental factors, as the temperature [43]. Once the nymphs eclose from the eggs, they undergo a series of moulds to finally mould into a terrestrial subimago that leaves the water. Strikingly, this sexually immature individual has to mould once more to become an imago, which is a singularity that only occurs in mayflies in contrast to all other insects that do not mould once they reach the adult stage [44, 45]. The mating occurs in flying swarms formed by hundreds of individuals several meters above the ground/water surface level [46–48].

**Figure 1.**
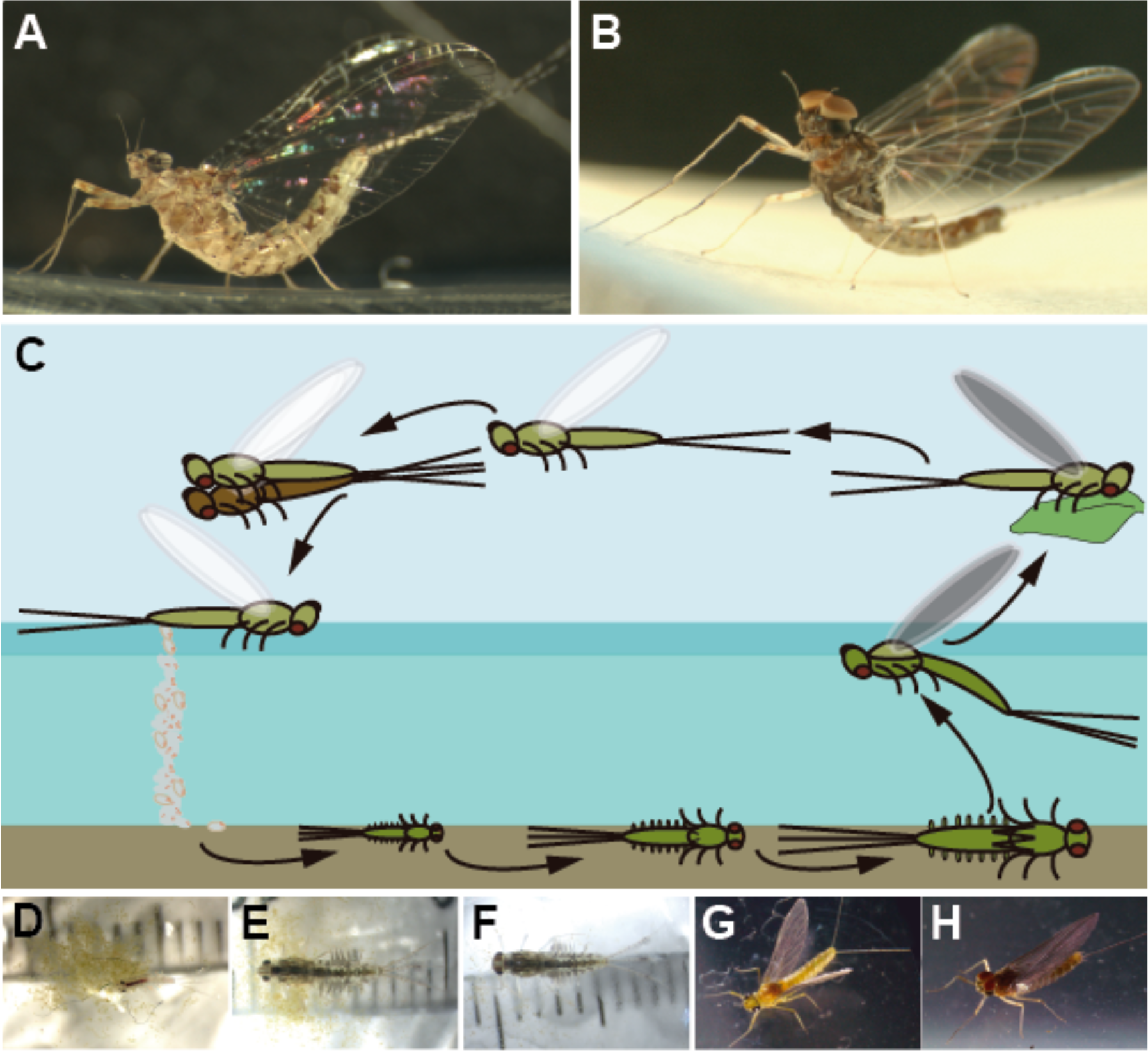
*C. dipterum* life cycle. **(A)** *C. dipterum* adult female **(B)** *C. dipterum* adult male. **(C)** Cartoon depicting *C. dipterum* life cycle. Female lays the eggs in a water stream where they hatch as juvenile nymphs. After several moults nymphs emerge from the water to the land as inmature subimagos. Then, they moult again to become sexually mature individuals that fly forming swarms to mate. **(D)** Early-mid nymph. **(E)** Late female nymph. **(F)** Late male nymph. **(G)** Female subimago. **(H)** Male subimago.

In most phylogenetic analyses, Ephemeropterans are grouped together with Odonata (damselflies and dragonflies) as the sister group of Neoptera, the rest of winged insects ([49] and references therein). Therefore, the extinct relatives of mayflies were the first insects developing wings, which makes extant mayflies a key organism to test the hypotheses postulated for the wing origin in pterygote insects. Their position in the phylogenetic tree also makes them an essential group to investigate segmentation, head specification and other morphogenetic processes occurring in the embryo, beyond the classical insect models already used to address these problems (*Drosophila, Tribolium, Oncopeltus*). Moreover, their particular life cycle with an aquatic and a terrestrial period makes mayflies a relevant organism to examine different adaptations to the land, such as the evolution of the extra-embryonic layers and other metabolic and physiologic traits derived from this complex life cycle, such as the hormonal control, ecdysis and metabolic rates.

By having established *C. dipterum* in the laboratory, we have now unlimited access to all embryonic and postembryonic stages through the year, permitting the study of fundamental processes that originated for the first time in Ephemeroptera or that are specific to the extant members of this order. Moreover, the development of a series of genomic and transcriptomic resources will facilitate comparative analyses at the genome, transcriptome and epigenomic levels that can clarify the role of certain genes and regulatory networks in the origin of those novelties. Finally, the highly regenerative capabilities of *C. dipterum*, together with its short life cycle (which lasts from forty to sixty days on average), make this species a significant and very useful system to investigate the regeneration of non-embryonic tissues in insects.

### *C. dipterum* continuous culture in the laboratory

*C. dipterum*, from the Baetidae family, is one of the few ovoviviparous ephemeropteran species: the female keeps the fertilized eggs inside the abdomen and only when they are ready to hatch, after ten to twenty days, the female sets down onto the surface of a water stream or pond and lays the eggs, that sink to the bottom ready to eclose. Just few seconds after the eggs are laid, the nymphs hatch [50].

Individual lines were established starting from single gravid females captured in Dos Hermanas (Sevilla, Spain) and Alfacar (Granada, Spain). In the laboratory, gravid females are kept in a petri dish with a wet filter paper to avoid its desiccation. After thirteen days, the female is placed on the surface of unchlorinated water in a beaker to let it lay the eggs. Usually, they immediately spawn if the eggs are ready to hatch, but the duration of embryogenesis is a bit variable, between thirteen and seventeen days. Thus, it is important to take the female back to the petri dish in case it did not spawn within the first minute on the water to avoid the laying of underdeveloped eggs. It is therefore advisable to try to induce the spawning during several days until reaching the appropriate moment when the embryos are fully developed. The amount of eggs a female can lay depends mainly on its nutritional condition. In general, bigger females produce larger clutches. In the laboratory, the females tend to lay between one hundred and three hundred eggs per clutch. Shortly after delivering the eggs, the females die.

The hatchlings take only a few seconds to hatch (Figure 2B-E, supplementary movie 1) as swimming nymphs. They instantly start feeding from algae that are placed at the bottom of the beaker. In the moment of hatching, the nymphs do not have external gills, it is only two moults later, approximately 72 hours after hatching that seven pairs of gills are visible in the first seven abdominal segments. The nymphs are kept in the unchlorinated water in the beaker during the whole juvenile period. A portion of the water is replaced once a week, though the frequency can be increased if the culture becomes cloudy due to an excess of mayfly faeces or overgrown of the algae. To improve the exchange of oxygen between the water and the air, a bubbling tube connected to an air pump is introduced in the water (Figure 2F). The nymphs feed regularly with *Chara*, filamentous algae, pulverised vegetarian fish flakes or small pieces of carrot that are added to the water.

**Figure 2.**
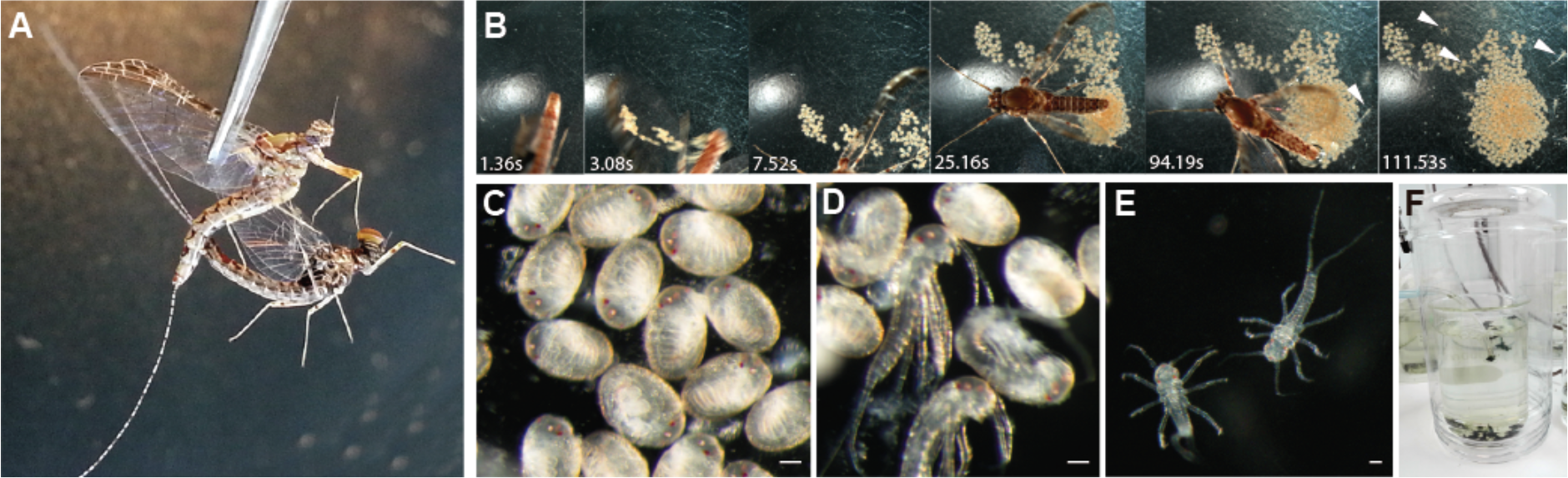
*C. dipterum* culture in the laboratory. **(A)** Couple of adults mating through forced copula. **(B)** Temporal sequence of a gravid female laying fertilised eggs that after 94 seconds hatch as swimming nymphs (white arrowheads). **(C-D)** Fertilised eggs and nymphs hatching. **(E)** Freshly hatched nymphs. **(F)** Culture system in the laboratory. Nymphs are in the beaker with bubbling water and algae. The beaker is placed inside a PET bottle to keep the subimagos once they emerge from the water. Scale bars: 50 μm

The beakers are placed inside PET bottles, so when nymphs reach the last nymphal stage and mould into a subimago that emerges and leaves the water, they can be easily recovered, as they cannot fly away and stay inside the bottles. To avoid water condensation that could damage the newly emerged subimagos while they stay inside the PET bottles, the plastic surface of their upper side is replaced by small net. The subimagos are carefully collected and kept for 24 hours in a tube with some wet paper to maintain the humidity and promote the last moult to imago, which happens some hours after the previous moult. To close the cycle in the laboratory, it is necessary to perform forced copulas [51], since mayflies mate during flight in large swarms [46–48]. To perform these matings, both, male and female are grasped very carefully by the wings with forceps. The female is placed with the ventral side upwards and the most posterior region of the male is brought close to the female seventh abdominal segment. Males, then, clasp the abdomen of the female using their genital forceps or stylus, allowing the contact of the two external genitalia to engage the copula (Fig. 2A). Copulas have a variable duration; they can last from few seconds to several minutes. During this time, males bend themselves to favour the fertilisation of the eggs. After the copula, the male is discarded and the female is kept in a petri dish with a small piece of humidified filter paper. The culture is maintained in a room at a constant temperature of 23 degrees Celsius and a 12:12 light:dark illumination cycle.

### *C. dipterum* embryogenesis

The establishment of *C. dipterum* culture in the laboratory allows the study of the complete embryogenesis of these mayflies by obtaining the embryos directly from the abdomen of gravid females. Once the embryos are collected from the female abdomen, it is possible to use antibodies and other markers to visualise the morphology of the embryo and morphogenetic processes occurring at specific developmental stages (Figure 3).

*C. dipterum* embryogenesis takes between thirteen and seventeen days, depending on the temperature. The morphogenesis in this species is similar to the previously described embryogenesis of other mayflies [52, 53]. Briefly, after egg cleavage the blastoderm is formed. Within the blastoderm, two populations of cells are soon distinguishable, the most posterior ones, that will form the germ disc and the larger and most anterior cells that will become the serosa (Figure 3A-B). Thereafter, the germ disc starts elongating and the future cephalic region and future caudal segment addition zone become apparent. During the following highly proliferative stages, as showed by an increased density of PH3-positive mitotic cells, especially in the most posterior regions (Figure C′-E′), the embryo elongates within the egg, adopting a S-shape (Figure 3C). The elongating embryo folds its most posterior region, which will correspond to abdominal segments, several times. After this phase, the developing *C. dipterum* reaches its final length and its segmentation starts. Segmentation happens from anterior to posterior, thus cephalic and thoracic appendages are the first to become visible (Figure 3D). Afterwards, the embryo undergoes a series of final developmental events in which the final form of the embryo is completed, such as the appearance of the caudal filament, two posterior cerci and the three ocelli and compound eyes (Figure 3E-F).

**Figure 3.**
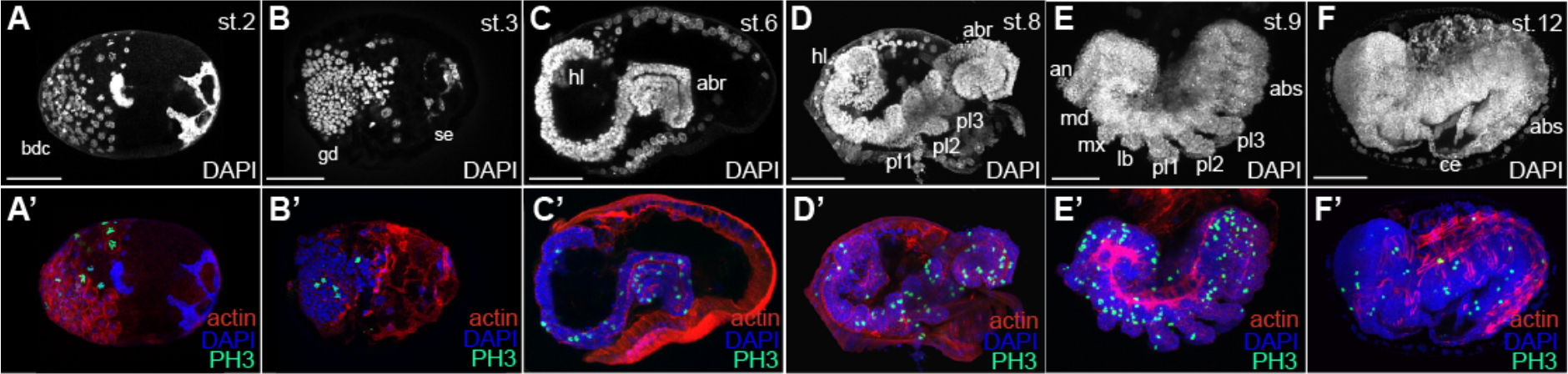
Representative phases of *C. dipterum* embryogenesis. Upper panels show embryo morphology detectable through DAPI staining (white). Lower panels show DAPI (nuclei, blue), Actin (cell contour cRed) and mitosis (anti-PH3, green). **(A-A′)** Blastoderm formation (stage 2: st. 2) bcd: blastoderm cells are replicating very actively, shown by PH3 staining **(A′)**, in green. **(B-B′)** Germ disc (gd) formation (st. 3). **(C-C′)** S-shaped embryo (st. 6). The germ band elongates backwards through active cell proliferation, mainly in the posterior region of the embryo (abr: abdominal region; hl: head lobe). **(D-D′)** Segmentation of the embryo (st. 8) starts from the cephalic (hl) and thoracic regions, which segments are already visible, towards the abdominal regions (abr). **(E-E′)** Proctodaeum formation (st. 9). Segmentation progresses, appendages enlarge and get segmented (an: antenna, md: mandible, mx: maxilla, lb: labium, pl: pro-leg). **(F-F′)** The abdominal regions are already segmented (abs: abdominal segments). Cercei (ce) are already visible. Dorsal closure proceeds. Scale bars: 50 μm

Beyond general morphology, access to all developmental stages allows the study of the development of specific tissues and organs. Using antibodies to detect neural structures, such as acetylated Tubulin (acTub, Figure 4) or probes against genes responsible for the patterning of specific regions of the nervous system, like *orthodenticle (otd)* or *engrailed (en)* to perform in situ hybridizations (Figure 5C-D), it is feasible to investigate the development of the nervous system, -or any other chosen embryonic territory-in embryos of the sister group to all other winged insects. Therefore, applying such techniques in *C. dipterum* embryos brings the possibility of studying fundamental processes (nervous system development, dorso-ventral patterning, segmentation, head development, extra-embryonic tissue specification, etc.) of insect development in an organism in a key position in the insect phylogeny.

**Figure 4.**
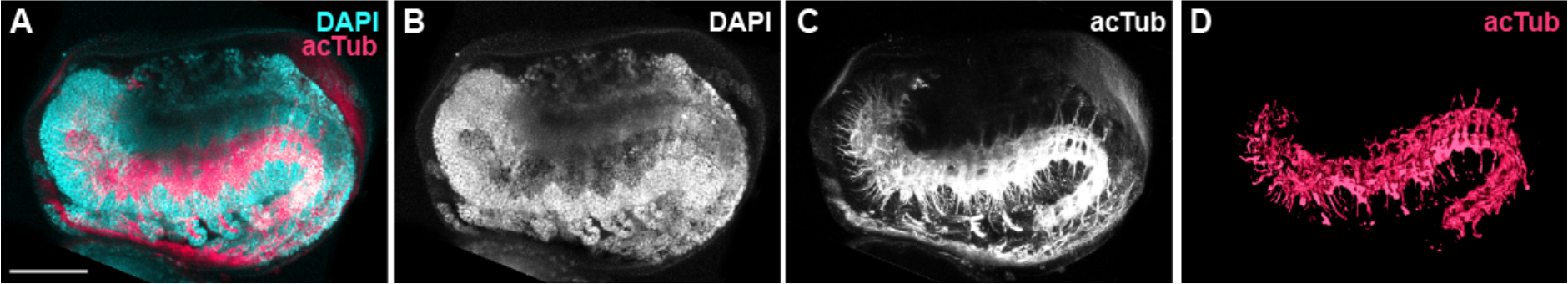
*C. dipterum* embryonic nervous system. **(A-C)** Embryo (DAPI staining reveals embryo morphology, B) exhibiting the ventral nervous cord (staining using anti-acetylated alphaTubulin antibody, C). **(D)** Surface reconstruction of the ventral nervous cord and its projections towards the appendages. Scale bars: 50 μm.

### The transcriptome of *C. dipterum*

Until now, there were no appropriate genomic tools available to investigate *C.dipterum* at the genetic level. Only a genomic survey sequencing for molecular markers to study *C. dipterum* population structure using 454 technology at low coverage has been reported [40, 54]. Therefore, to explore *C. dipterum* gene content, we sequenced the transcriptome of a male nymph. The assembly of the paired-end reads, using Trinity RNA-Seq *de novo* assembly software [55] resulted in 117233 transcripts. From these 117233 transcripts we obtained 95053 peptide sequences using transDecoder software [56] to get the longest translated ORFs. Running BLASTp, we got a list of 15799 sequences from UniRef90 database which showed homology to other sequences (e-value < 10e-6). These hits showed a majority of results, more than 80% (13059 best hits), within the hexapoda (insects and Collembola). The second most frequent groups of hits fell within Arthropoda (Chelicerata and Myriapoda, 4,24%) and Crustacea (4,15%) categories, which demonstrated the good quality of the assembly (Figure 5B). Less frequent categories present in our best hit results corresponded, on the one hand, to Bacteria and virus which probably derive from the mayfly microbiota and on the other hand, Plantae and Red algae, which most likely belonged to the gut content of the specimen at the moment of the RNA extraction, as *C. dipterum* feeds on algae and plants.

**Figure 5.**
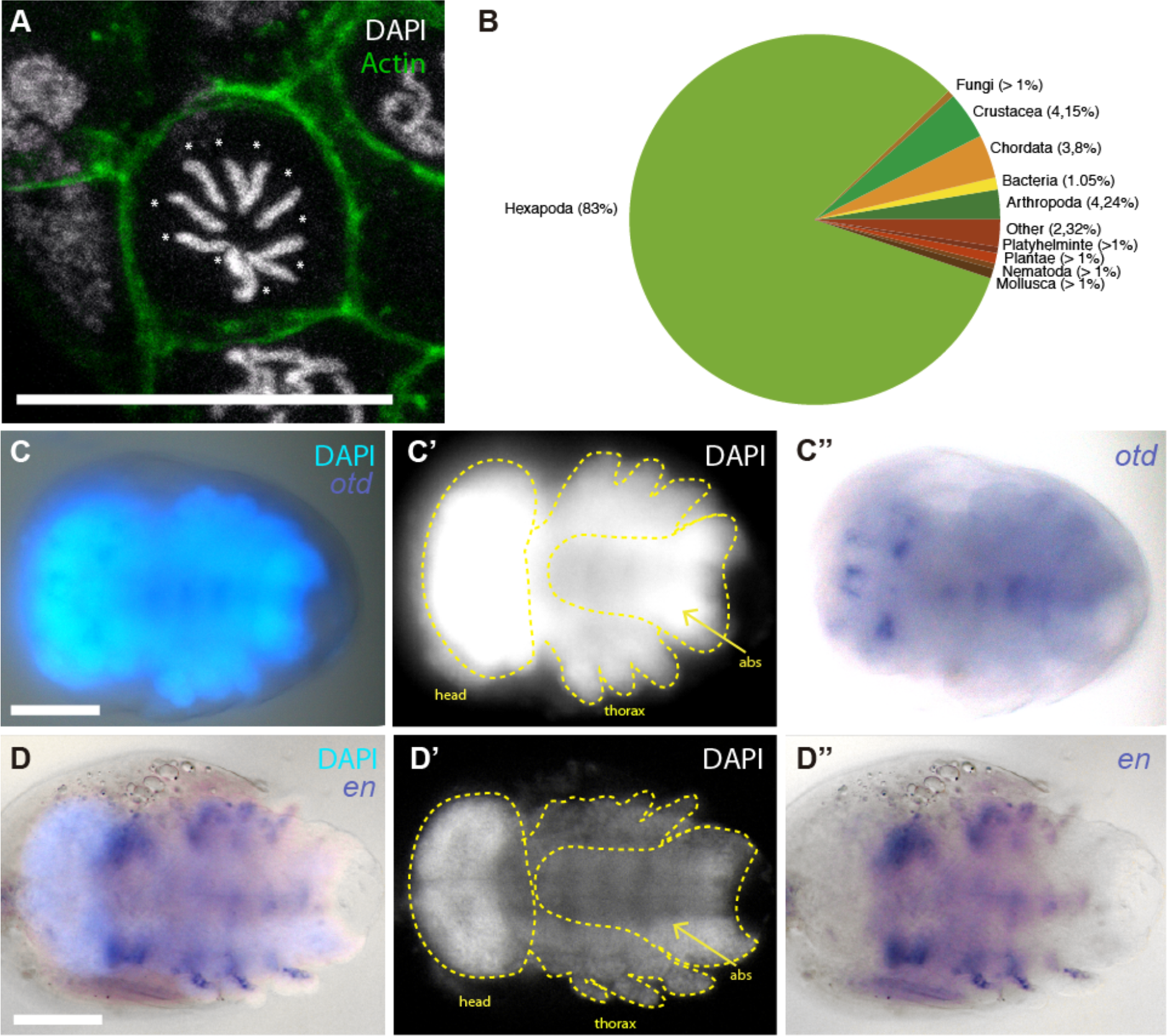
Genomics and transcriptomic tools. **(A)** The genome of *C. dipterum* is structured in a karyotype of 2n = 10 [97, 98]. Somatic embryonic cell showing condensed DNA in chromosomes (each of them highlighted with an asterisk). DNA stained with DAPI (white) and cell membrane visible through Phalloidin-Rhodamin staining (green). Scale bar: 20 μm **(B**) Pie chart representing top blastp results of unigenes against UniRef90 protein database. **(C-C′ ′)** *otd* expression pattern **(C, C′ ′)** in the mayfly embryo visible through DAPI staining **(C, C′)**. **(D-D′′)** *en* expression domain (D, D′ ′) in *C. dipterum* embryo **(D, D′)**.

Although the transcriptome generated was obtained from a single male nymph, and it thus represents only the genes that are expressed at that particular developmental stage, it can nevertheless serve as a very useful resource to identify homologous transcripts and to design probes to perform subsequent expression pattern analyses of genes of interest expressed during nymphal stages and other stages, as in the case of *orthodenticle* (*otd*) and *engrailed* (*en*) (real time PCRs, in situ hybridization, Figure 5C-D).

This transcriptome assembly is a first step in order to have resources that can be used to tackle questions in the evolution of first winged insects at a genomic/transcriptomic level. Nevertheless, more tools are needed, so the high quality genome sequencing project that is currently in progress will provide an invaluable resource and a platform for subsequent analyses (ATAC-Seq, Chip-Seq, etc.) to investigate long standing questions related to the origin of pterygotes and other important traits that contributed to the diversification of insects.

### The regenerative potential of *C. dipterum*

The capacity to regenerate lost or damaged organs, body parts (or even whole organisms) is widespread through the animal kingdom [57–59]. Several phyla, such as Cnidarian, Platyhelminthes, annelids, arthropods or vertebrates, have this ability that can vary in several aspects. Different species or even phyla have very different regenerative capabilities; for instance, Platyhelminthes (flatworms) are able to regenerate a complete organism from few hundreds of cells [60–66] whereas mammals have lost most of their regenerative potential, and they are able to regenerate only particular tissues or organs in specific conditions [67, 68]. The mechanisms used in the regeneration process are also different depending on the species; flatworms use totipotent cells, the neoblasts, to regenerate, while other organisms, such as the crustacean *Parhyale*, rely on the dedifferentiation of cell populations to re-grow an amputated limb [58, 69, 70]. Despite the diversity of species that are able to regenerate and the varying modes, mechanisms and degrees of their regeneration capabilities, only a small number of organisms have been used to investigate how regeneration occurs. Thus, increasing the diversity of regenerating model species will greatly contribute to study whether general rules work in the morphogenetic process or whether the same gene networks and regulatory modules are common to different phyla with regeneration potential. This is particular evident for insects, where only a handful of species have been used as models for regeneration studies, *Drosophila melanogaster* [71–75], cockroaches [76, 77] and *Gryllus bimaculatus* [77–79]. The genetic toolkit available for *Drosophila* manipulation has greatly contributed to the identification of gene networks involved in the regeneration event [80–86]. However, the regenerative potential of *Drosophila* is quite limited, thus, it has been only described in imaginal discs, which are highly proliferative and undifferentiated organs that will give rise to the adult appendages, and in the gut, which, in the same manner as the imaginal discs, possesses a high proliferative population of cells, including intestinal stem cells that promote the regrowth of the gut after damage [75, 87]. On the other hand, crickets and cockroaches have been developed as models for limb regeneration, giving the possibility of studying fully functional organs with terminally differentiated cell types [77, 78, 88–92]. However, crickets take one month to complete the regeneration of a leg, while in cockroaches the whole process can last more than eighteen weeks [89].

Dewitz, already in 1890, described that mayfly nymphs were able to regenerate their gills completely after amputation [93]. Since then, several researchers [94, 95] confirmed these observations. Indeed, we observed that *C. dipterum* is able to regenerate gills, antennae, cerci and legs completely in a very short period of time, ranging from six to nine days (Fig. 6). For instance, after amputation of the 3rd pair of legs, *C. dipterum* takes less than 24 hours to heal the wound and only 72 hours to exhibit a clear re-growth of the appendage, completing the entire process in a period of no more than seven days (Fig. 6). Thus, *C. dipterum* has extraordinarily rapid regenerative capabilities, that could give this species a privileged status because of its fast regeneration of postembryonic, fully functional organs.

**Figure 6.**
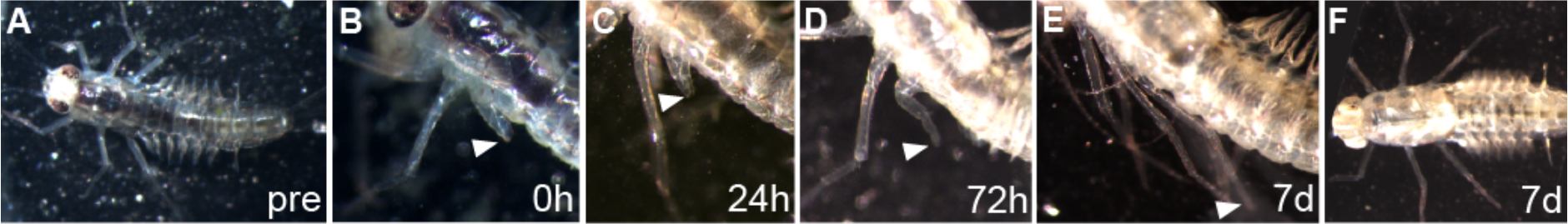
Leg regeneration of a *C. dipterum* nymph. **(A)** Mayfly nymph before amputation of the 3rd leg. **(B)** Nymphal leg (white arrowhead) immediately after amputation. **(C)** 24 hours after amputation the wound is healed (arrowhead). **(D)** 72 hours after amputation, the tissue is already partially regenerated. **(E-F)** After seven days, the amputated leg has recovered its initial size and shape with all the segments perfectly formed.

## Conclusions

Although Ephemeropterans have been the focus of biogeography, taxonomy and ecology studies, until now they have been very rarely used as a laboratory model to address developmental and evolutionary questions [34, 96], despite the fact they are fundamental to understand insect evolution at multiple time scales. Here we present *C. dipterum* as an emergent model for evo-devo studies. There are several traits in this species that make it especially helpful to answer long-standing questions in evolutionary biology and its establishment in the laboratory provides great advantages. First, the setting up of a continuous culture system in the laboratory facilitates the access to all the developmental stages, and as for the particular case of the ovoviviparism of *C. dipterum*, it allows having high number of synchronised embryos. The continuous culture also permits to obtain large amounts of material that can be used in genomics and transcriptomics assays. Second, the use of forced copulas ensures a complete control on the matings, so it is feasible to have inbred lines to reduce genetic heterozygosity to perform genetic experiments and when applying functional genomics techniques. Moreover, the relatively short life cycle of *C.dipterum* permits the investigation of embryonic and postembryonic processes in a brief period of time and making experimental designs feasible. Although RNA interference techniques are not established in the mayfly yet, its aquatic life phase allows us to perform drug treatments, by just adding the molecule to the water, to interfere with signalling pathways in order to investigate their function in specific conditions. However, these functional experiments are limited by the amount of drugs available to alter specific gene networks, thus, efforts must be made to set up interference RNA and CRISPR/Cas9 methods to downregulate genes or edit the genome in *C. dipterum.*

Moreover, the generation of -omic resources, through the sequencing of *C. dipterum* genome and different tissue and stage specific RNA-seq datasets, and the development of protocols to investigate changes in regulatory regions of the genome, such as ATAC-seq and ChIP-seq techniques, will provide a great resource to investigate evolutionary and developmental questions at the genomic level.

Beyond the technical and methodological advantages that *C. dipterum* system confers, its key phylogenetic position, ecology, physiology, and plasticity make mayflies an essential order to investigate very diverse topics, from genomic and morphogenetic events that occurred at the origin of winged insects, the origin of metamorphosis and hormone control of ecdysis, to the regenerative potential in insects.

## Acknowledgements

We thank David Funk, Pablo Jáimez and Julio Miguel Luzón for helping and advise on the collection and establishment of the *C. dipterum* continuous system in the laboratory and Zeynep Kalender for helping on the initial analyses of the transcriptome. We also thank the CABD Fish facility and members of the Martinez-Morales lab for providing filamentous algae to feed the mayflies, the CABD ALMI platform for confocal microscopy support and Ignacio Maeso for critically reading the manuscript. This project has been funded by the European Union’s Horizon 2020 research and innovation programme under the Marie Sklodowska-Curie grant agreement 657732 to IA and *Apoyo a Unidades de Excelencia Maía de Maeztu* (MDM-2016-0687) and MINECO (Spain) through grant no. BFU2015-66040-P to FC from the Ministry of Economy, Industry and Competitiveness of Spain.

## Authors’ contributions

IA and FC conceived and designed the study. IA and CM-B established and maintained *C. dipterum* culture. IA and CM-B performed embryogenesis characterisation. IA, IMG-F and AL-C performed in situ hybridization experiments. IA, KD and SA generated and analysed the transcriptome data. CM-B and IA performed regeneration assays. IA prepared the figures and wrote the manuscript with the help of FC and inputs from all authors. All authors read, corrected and approved the final manuscript.

## References

1. Misof B, Liu S, Meusemann K, Peters RS, Donath A, Mayer C, Frandsen PB, Ware J, Flouri T, Beutel RG et al: Phylogenomics resolves the timing and pattern of insect evolution. Science 2014, 346(6210):763–767.

2. Collins NM, Thomas JA: The conservation of insects and their habitats. Academic Press, London 1991.

3. Lozano-Fernandez J, Carton R, Tanner AR, Puttick MN, Blaxter M, Vinther J, Olesen J, Giribet G, Edgecombe GD, Pisani D: A molecular palaeobiological exploration of arthropod terrestrialization. Philos Trans R Soc Lond B Biol Sci 2016, 371(1699).

4. Rota-Stabelli O, Daley Allison C, Pisani D: Molecular Timetrees Reveal a Cambrian Colonization of Land and a New Scenario for Ecdysozoan Evolution. Current Biology 2013, 23(5):392–398.

5. Little C: The colonisation of land: origins and adaptations of terrestrial animals. Cambridge University Press, Cambridge [Cambridgeshire]; New York 1983:241–272.

6. Hilbrant M, Almudi I, Leite DJ, Kuncheria L, Posnien N, Nunes MD, McGregor AP: Sexual dimorphism and natural variation within and among species in the Drosophila retinal mosaic. BMC Evol Biol 2014, 14:240.

7. Kittelmann S, Buffry AD, Franke FA, Almudi I, Yoth M, Sabaris G, Couso JP, Nunes MDS, Frankel N, Gomez-Skarmeta JL et al: Gene regulatory network architecture in different developmental contexts influences the genetic basis of morphological evolution. PLoS Genet 2018, 14(5):e1007375.

8. Smith SJ, Rebeiz M, Davidson L: From pattern to process: studies at the interface of gene regulatory networks, morphogenesis, and evolution. Current Opinion in Genetics & Development 2018, 51:103–110.

9. Schmidt-Ott U, Lynch JA: Emerging developmental genetic model systems in holometabolous insects. Current Opinion in Genetics & Development 2016, 39:116–128.

10. Chipman AD: Oncopeltus fasciatus as an evo-devo research organism. genesis 2017, 55(5):e23020.

11. Korb J, Belles X: Juvenile hormone and hemimetabolan eusociality: a comparison of cockroaches with termites. Current Opinion in InsectScience 2017, 22:109–116.

12. Santos ME, Le Bouquin A, Crumière AJJ, Khila A: Taxon-restricted genes at the origin of a novel trait allowing access to a new environment. Science 2017, 358(6361):386–390.

13. Toubiana W, Khila A: The benefits of expanding studies of trait exaggeration to hemimetabolous insects and beyond morphology. Current Opinion in Genetics & Development 2016, 39:14–20.

14. Schmidt-Ott U, Kwan CW: Morphogenetic functions of extraembryonic membranes in insects. Current Opinion in Insect Science 2016, 13:86–92.

15. Jacobs CGC, Rezende GL, Lamers GEM, van der Zee M: The extraembryonic serosa protects the insect egg against desiccation. Proceedings of the Royal Society B: Biological Sciences 2013, 280(1764).

16. van der Zee M, Berns N, Roth S: Distinct Functions of the *Triboliumzerknüllt* Genes in Serosa Specification and Dorsal Closure. Current Biology 2005, 15(7):624–636.

17. Jacobs CGC, Spaink HP, van der Zee M: The extraembryonic serosa is afrontier epithelium providing the insect egg with a full-range innateimmune response. eLife 2014, 3:e04111.

18. Hilbrant M, Horn T, Koelzer S, Panfilio KA: The beetle amnion and serosa functionally interact as apposed epithelia. eLife 2016, 5:e13834.

19. Auman T, Chipman AD: Growth zone segmentation in the milkweed bug Oncopeltus fasciatus sheds light on the evolution of insect segmentation. BMC Evolutionary Biology 2018, 18(1):178.

20. Zhu X, Rudolf H, Healey L, François P, Brown SJ, Klingler M, El-Sherif E:Speed regulation of genetic cascades allows for evolvability in the body plan specification of insects. Proceedings of the National Academy of Sciences 2017, 114(41):E8646–E8655.

21. Liu PZ, Kaufman TC: Short and long germ segmentation: unanswered questions in the evolution of a developmental mode. Evolution & Development 2005, 7(6):629–646.

22. Stahi R, Chipman AD: Blastoderm segmentation in Oncopeltus fasciatus and the evolution of insect segmentation mechanisms. Proceedings of the Royal Society B: Biological Sciences 2016, 283(1840).

23. Peel A: The evolution of arthropod segmentation mechanisms. BioEssays 2004, 26(10):1108–1116.

24. Magri MS, Domínguez-Cejudo MA, Casares F: Wnt controls the medial-lateral subdivision of the *Drosophila* head. Biology Letters 2018, 14(7).

25. Posnien N, Schinko JB, Kittelmann S, Bucher G: Genetics, development and composition of the insect head – A beetle’s view. Arthropod Structure & Development 2010, 39(6):399–410.

26. Hamilton KGA: The insect wing. Part 1. Origin and development of wings from notal lobes. 1971, 44:421–433. J Kansas Entomol Soc 1971, 44:421–433.

27. Rasnitsyn AP: A modified paranotal theory of insect wing origin. Journal of Morphology 1981, 168(3):331–338.

28. Ohde T, Yaginuma T, Niimi T: Insect Morphological Diversification Through the Modification of Wing Serial Homologs. Science 2013, 340(6131):495–498.

29. Averof M, Cohen SM: Evolutionary origin of insect wings from ancestral gills. Nature 1997, 385(6617):627–630.

30. Clark-Hachtel CM, Tomoyasu Y: Exploring the origin of insect wings from an evo-devo perspective. Current Opinion in Insect Science 2016, 13:77–85.

31. Kukalova-Peck J: Origin and evolution of insect wings and their relation to metamorphosis, as documented by the fossil record. Journal of Morphology 1978, 156(1):53–125.

32. Kukalová-Peck J: Origin of the insect wing and wing articulation from the arthropodan leg. Canadian Journal of Zoology 1983, 61(7):1618–1669.

33. Linz DM, Tomoyasu Y: Dual evolutionary origin of insect wings supported by an investigation of the abdominal wing serial homologs in Tribolium. Proc Natl Acad Sci U S A 2018, 115(4):E658–E667.

34. Niwa N, Akimoto-Kato A, Niimi T, Tojo K, Machida R, Hayashi S:Evolutionary origin of the insect wing via integration of two developmental modules. Evolution & Development 2010, 12(2):168–176.

35. Barber-James HM, Gattolliat J-L, Sartori M, Hubbard MD: Global diversity of mayflies (Ephemeroptera, Insecta) in freshwater. Hydrobiologia 2008,595(1):339–350.

36. Sartori M, Brittain JE: Chapter 34 - Order Ephemeroptera. In: Thorp and Covich’s Freshwater Invertebrates (Fourth Edition). Edited by Thorp JH, Rogers DC. Boston: Academic Press; 2015: 873–891.

37. Chou H, Pathmasiri W, Deese-spruill J, Sumner SJ, Jima DD, Funk DH, Jackson JK, Sweeney BW, Buchwalter DB: The Good, the Bad, and the Lethal: Gene Expression and Metabolomics Reveal Physiological Mechanisms Underlying Chronic Thermal Effects in Mayfly Larvae (Neocloeon triangulifer). Frontiers in Ecology and Evolution 2018, 6(27).

38. Peschke K, Geburzi J, Köhler H-R, Wurm K, Triebskorn R: Invertebrates as indicators for chemical stress in sewage-influenced stream systems:Toxic and endocrine effects in gammarids and reactions at the community level in two tributaries of Lake Constance, Schussen and Argen. Ecotoxicology and Environmental Safety 2014, 106:115–125.

39. Scarduelli L, Giacchini R, Parenti P, Migliorati S, Di Brisco AM, Vighi M: Natural variability of biochemical biomarkers in the macro-zoobenthos:Dependence on life stage and environmental factors. Environmental Toxicology and Chemistry 2017, 36(11):3158–3167.

40. Rutschmann S, Detering H, Simon S, Funk DH, Gattolliat J-L, Hughes SJ, Raposeiro PM, DeSalle R, Sartori M, Monaghan MT: Colonization and diversification of aquatic insects on three Macaronesian archipelagos using 59 nuclear loci derived from a draft genome. Molecular Phylogenetics and Evolution 2017, 107:27–38.

41. Vuataz L, Rutschmann S, Monaghan MT, Sartori M: Molecular phylogeny and timing of diversification in Alpine Rhithrogena (Ephemeroptera: Heptageniidae). BMC Evolutionary Biology 2016, 16(1):194.

42. Gueuning M, Suchan T, Rutschmann S, Gattolliat J-L, Jamsari J, Kamil AI, Pitteloud C, Buerki S, Balke M, Sartori M et al: Elevation in tropical sky islands as the common driver in structuring genes and communities of freshwater organisms. Scientific Reports 2017, 7(1):16089.

43. Humpesch UH: Effect of fluctuating temperature on the duration of embryonic development in two Ecdyonurus spp. and Rhithrogena cf. hybrida (Ephemeroptera) from Austrian streams. Oecologia 1982, 55(3):285–288.

44. Edmunds JGF, McCafferty WP: The mayfly subimago. Ann Rev Entomology 1988, 33: 509–29.

45. Maiorana VC: Why do adult insects not moult? Biological Journal of the Linnean Society 1979, 11(3):253–258.

46. Harker JE: Swarm behaviour and mate competition in mayflies (Ephemeroptera). Journal of Zoology 1992, 228(4):571–587.

47. Allan JD, Flecker AS: The mating biology of a mass-swarming mayfly. Animal Behaviour 1989, 37:361–371.

48. Peckarsky BL, McIntosh AR, Caudill CC, Dahl J: Swarming and mating behavior of a mayfly Baetis bicaudatus suggest stabilizing selection for male body size. Behavioral Ecology and Sociobiology 2002, 51(6):530–537.

49. Simon S, Blanke A, Meusemann K: Reanalyzing the Palaeoptera problem - The origin of insect flight remains obscure. Arthropod Structure & Development 2018, 47(4):328–338.

50. Berner L: Ovoviviparous Mayflies in Florida. The Florida Entomologist 1941, 24(2):32–34.

51. Huff BL, Jr.; McCafferty, W. P.: Parthenogenesis and experimental reproductive biology in four species of the mayfly genus stenonema. The Wasmann Journal of Biology 1974, 32(2).

52. Bohle HW: The effect of temperature on embryogenesis and diapause of Ephemerella ignita (Poda). Oecologia 1972, 10(3):253–268.

53. Tojo K, Machida R: Early embryonic development of the mayfly Ephemera japonica McLachlan (Insecta: Ephemeroptera, Ephemeridae). J Morphol 1998, 238(3):327–335.

54. Rutschmann S, Detering H, Simon S, Fredslund J, Monaghan MT:discomark: nuclear marker discovery from orthologous sequences using draft genome data. Molecular Ecology Resources 2017, 17(2):257–266.

55. Grabherr MG, Haas BJ, Yassour M, Levin JZ, Thompson DA, Amit I, Adiconis X, Fan L, Raychowdhury R, Zeng Q et al: Full-length transcriptome assembly from RNA-Seq data without a reference genome. Nat Biotechnol 2011, 29(7):644–652.

56. Haas BJ, Papanicolaou A, Yassour M, Grabherr M, Blood PD, Bowden J, Couger MB, Eccles D, Li B, Lieber M et al: De novo transcript sequence reconstruction from RNA-seq using the Trinity platform for reference generation and analysis. Nature protocols 2013, 8(8):1494–1512.

57. Bely AE, Nyberg KG: Evolution of animal regeneration: re-emergence of afield. Trends in Ecology & Evolution 2010, 25(3): 161–170.

58. Grillo M, Konstantinides N, Averof M: Old questions, new models:unraveling complex organ regeneration with new experimental approaches. Curr Opin Genet Dev 2016, 40:23–31.

59. Morgan TH: Regeneration. The Macmillan Company 1901.

60. Atabay KD, LoCascio SA, de Hoog T, Reddien PW: Self-organization and progenitor targeting generate stable patterns in planarian regeneration. Science 2018, 360(6387):404–409.

61. Cebria F, Adell T, Salo E: Rebuilding a planarian: from early signaling to final shape. Int J Dev Biol 2018, 62(6-7-8):537–550.

62. Egger B, Gschwentner R, Hess MW, Nimeth KT, Adamski Z, Willems M, Rieger R, Salvenmoser W: The caudal regeneration blastema is an accumulation of rapidly proliferating stem cells in the flatworm Macrostomum lignano. BMC Dev Biol 2009, 9:41.

63. Egger B, Ladurner P, Nimeth K, Gschwentner R, Rieger R: The regeneration capacity of the flatworm Macrostomum lignano--on repeated regeneration, rejuvenation, and the minimal size needed forregeneration. Dev Genes Evol 2006, 216(10):565–577.

64. Fincher CT, Wurtzel O, de Hoog T, Kravarik KM, Reddien PW: Cell type transcriptome atlas for the planarian Schmidtea mediterranea. Science 2018, 360(6391).

65. Nimeth KT, Egger B, Rieger R, Salvenmoser W, Peter R, Gschwentner R:Regeneration in Macrostomum lignano (Platyhelminthes): cellulardynamics in the neoblast stem cell system. Cell Tissue Res 2007,327(3):637–646.

66. Reddien PW: The Cellular and Molecular Basis for Planarian Regeneration. Cell 2018, 175(2):327–345.

67. Quaife-Ryan GA, Sim CB, Ziemann M, Kaspi A, Rafehi H, Ramialison M, El-Osta A, Hudson JE, Porrello ER: Multicellular Transcriptional Analysis of Mammalian Heart Regeneration. Circulation 2017, 136(12):1123–1139.

68. Zacharias WJ, Frank DB, Zepp JA, Morley MP, Alkhaleel FA, Kong J, Zhou S, Cantu E, Morrisey EE: Regeneration of the lung alveolus by an evolutionarily conserved epithelial progenitor. Nature 2018,555(7695):251–255.

69. Konstantinides N, Averof M: A Common Cellular Basis for Muscle Regeneration in Arthropods and Vertebrates. Science 2014,343(6172):788–791.

70. Alwes F, Enjolras C, Averof M: Live imaging reveals the progenitors andcell dynamics of limb regeneration. eLife 2016, 5:e19766.

71. Hariharan IK, Serras F: Imaginal disc regeneration takes flight. Curr OpinCell Biol 2017, 48:10–16.

72. Repiso A, Bergantiños C, Corominas M, Serras F: Tissue repair and regeneration in Drosophila imaginal discs. Development, Growth &Differentiation 2011, 53(2): 177–185.

73. Vizcaya-Molina E, Klein CC, Serras F, Mishra RK, Guigo R, Corominas M:Damage-responsive elements in Drosophila regeneration. Genome Res 2018.

74. Ahmed-de-Prado S, Baonza A: Drosophila as a Model System to Study Cell Signaling in Organ Regeneration. Biomed Res Int 2018, 2018:7359267.

75. Guo Z, Lucchetta E, Rafel N, Ohlstein B: Maintenance of the adult Drosophila intestine: all roads lead to homeostasis. Curr Opin Genet Dev 2016, 40:81–86.

76. French V: Leg regeneration in the cockroach, Blatellagermanica. II Regeneration from a non-congruent tibial graft/host junction 1976, 35(2):267–301.

77. Nakamura T, Mito T, Bando T, Ohuchi H, Noji S: Molecular and Cellular Basis of Regeneration and Tissue Repair. Cellular and Molecular LifeSciences 2007, 65(1):64.

78. Bando T, Mito T, Hamada Y, Ishimaru Y, Noji S, Ohuchi H: Molecular mechanisms of limb regeneration: insights from regenerating legs of the cricket Gryllus bimaculatus. Int J Dev Biol 2018, 62(6-7-8):559–569.

79. Das S: Morphological, Molecular, and Hormonal Basis of Limb Regeneration across Pancrustacea. Integrative and Comparative Biology 2015, 55(5):869–877.

80. Ahmed-de-Prado S, Diaz-Garcia S, Baonza A: JNK and JAK/STAT signalling are required for inducing loss of cell fate specification during imaginal wing discs regeneration in Drosophila melanogaster. Dev Biol 2018, 441 (1):31–41.

81. Bergantinos C, Corominas M, Serras F: Cell death-induced regeneration in wing imaginal discs requires JNK signalling. Development 2010,137(7): 1169–1179.

82. Blanco E, Ruiz-Romero M, Beltran S, Bosch M, Punset A, Serras F, Corominas M: Gene expression following induction of regeneration in Drosophila wing imaginal discs. Expression profile of regenerating wing discs. BMC Dev Biol 2010, 10:94.

83. Bosch M, Bishop SA, Baguna J, Couso JP: Leg regeneration in Drosophila abridges the normal developmental program. Int J Dev Biol 2010, 54(8-9):1241–1250.

84. Bosch M, Serras F, Martin-Blanco E, Baguna J: JNK signaling pathway required for wound healing in regenerating Drosophila wing imaginal discs. Dev Biol 2005, 280(1):73–86.

85. Harris RE, Setiawan L, Saul J, Hariharan IK: Localized epigenetic silencing of a damage-activated WNT enhancer limits regeneration in mature Drosophila imaginal discs. Elife 2016, 5.

86. Santabarbara-Ruiz P, Lopez-Santillan M, Martinez-Rodriguez I, Binagui-Casas A, Perez L, Milan M, Corominas M, Serras F: ROS-Induced JNK andp38 Signaling Is Required for Unpaired Cytokine Activation during Drosophila Regeneration. PLoS Genet 2015, 11(10):e1005595.

87. Schwartz S, Rhiner C: Reservoirs for repair? Damage-responsive stemcells and adult tissue regeneration in Drosophila. Int J Dev Biol 2018,62(6-7-8):465–471.

88. Bando T, Ishimaru Y, Kida T, Hamada Y, Matsuoka Y, Nakamura T, Ohuchi H, Noji S, Mito T: Analysis of RNA-Seq data reveals involvement of JAK/STAT signalling during leg regeneration in the cricket Gryllusbimaculatus. Development 2013, 140(5):959–964.

89. Hamada Y, Bando T, Nakamura T, Ishimaru Y, Mito T, Noji S, Tomioka K, Ohuchi H: Leg regeneration is epigenetically regulated by histone H3K27 methylation in the cricket Gryllus bimaculatus. Development 2015,142(17):2916–2927.

90. Hamada Y, Tokuoka A, Bando T, Ohuchi H, Tomioka K: Enhancer of zesteplays an important role in photoperiodic modulation of locomotor rhythm in the cricket, Gryllus bimaculatus. Zoological Lett 2016, 2:5.

91. Nakamura T, Mito T, Miyawaki K, Ohuchi H, Noji S: EGFR signaling isrequired for re-establishing the proximodistal axis during distal legregeneration in the cricket Gryllus bimaculatus nymph. Developmental Biology 2008, 319(1):46–55.

92. Tanaka A, Akahane H, Ban Y: The problem of the number of tarsomeres in the regenerated cockroach leg. Journal of Experimental Zoology 1992,262(1):61–70.

93. Dewitz H: Zool Anz. 13 1890, 525.

94. Cuénot L: L’adaptation: les Impr. réunies; 1925.

95. Wingfield CA: The Function of the Gills of Mayfly Nymphs from Different Habitats. Journal of Experimental Biology 1939, 16(3):363–373.

96. O’Donnell BC, Jockusch EL: The expression of wingless and Engrailed in developing embryos of the mayfly Ephoron leukon (Ephemeroptera:Polymitarcyidae). Development Genes and Evolution 2010, 220(1):11–24.

97. Kiauta B, M. Maw: Behaviour of the spermatocyte chromosomes of the mayfly, Cloeon dipterum (Linnaeus, 1761) s.1. (Ephemeroptera:Baetidae), with a note on the cytology of the order. Genen Phaenen 19(2/3): 31–39 1977.

98. Wolf E: Zur Karyologie der Eireifung und Furchung bei Cloeon dipterum L. (Bengtsson)(Ephemerida, Baetidae). Biol Zbl 79: 153–198 1960.

